# Bioplastic Production Potential of *Azospira suillum* PS: Growth-Associated PHB Production Under Aerobic and Anaerobic Conditions

**DOI:** 10.1101/2025.11.12.688118

**Authors:** David A.O. Meier, Benjamin Glazer, V. Celeste Lanclos, Hans K. Carlson, John D. Coates

## Abstract

Plastics are indispensable in modern society, yet less than 10% of the 380 million tons produced annually are recycled. This accumulation of non-degradable waste has intensified the search for sustainable alternatives. Polyhydroxyalkanoates (PHAs), particularly polyhydroxybutyrate (PHB), are biodegradable polymers produced by microorganisms. Here, we investigate the capacity of *Azospira suillum* PS, a model facultative perchlorate-reducing bacterium, to synthesize PHB. We show that unlike most organisms, PHB production is growth-associated both aerobically, and anaerobically with perchlorate as an electron acceptor under moderate nitrogen deplete conditions. Genomic analysis revealed four *phaC* homologs, with one, *phaC4*, co-localized with *phaB* and *phaR*, suggesting a conserved operon. While three of the *phaC* genes were successfully deleted, *phaC4* could not be disrupted, indicating an essential role for growth. Redox profiling revealed a positive correlation between PHB accumulation and NADPH/NADP^+^ ratios, supporting the role of cytoplasmic reducing equivalents in PHB synthesis. This study presents the first demonstration of PHA production under perchlorate-respiring conditions and one of the few documented cases of growth-associated PHA synthesis. It offers new insights into the genetic and metabolic basis of PHB production in *A. suillum* PS, underscoring its potential as a platform organism for sustainable, growth-coupled bioplastic production.

## Introduction

The use of processed polymers by humans dates to at least 1600 B.C.E. (Hosler et al., 1999), but the invention of modern thermoplastics by Charles Goodyear in 1839 revolutionized their industrial and environmental impact. Today, thermoplastics are indispensable across nearly every sector due to their lightweight, durable, and versatile properties (Andrady & Neal, 2009). This utility has driven global plastic production to over 380 million tons annually (Geyer et al., 2017). However, nearly all conventional plastics are derived from fossil hydrocarbons and are non-biodegradable, leading to long-term environmental accumulation. This persistent pollution, along with the carbon footprint associated with petrochemical feedstocks, has accelerated interest in sustainable alternatives such as bioplastics—materials synthesized from renewable sources.

Polyhydroxyalkanoates (PHAs) are one such family of bioplastics that are fully compostable, produced by microorganisms as intracellular carbon and energy storage polymers (Macrae & Wilkinson, 1958). PHAs are classified by monomer length into short-chain-length (scl-PHA, C3–C5) and medium-chain-length (mcl-PHA, C6–C14) types, with copolymers offering diverse material properties (Steinbüchel, 1991; Chen et al., 2015; McAdam et al., 2020). This chemical versatility allows tailoring of rigidity, flexibility, and degradability for various applications (Gao et al., 2011). However, commercial PHA adoption is constrained by cost of production—typically $3–8 per kg in 2025, compared to ∼$1.40 for petrochemical polyethylene—with global production representing less than 0.5% of total plastics (Rosenboom et al., 2022). Reducing cost of production through improvement of microbial titers and yields requires deeper insight into PHA biosynthetic regulation and physiology.

Among PHAs, polyhydroxybutyrate (PHB) is the best studied, first identified in *Bacillus subtilis* in 1926 (Lemoigne). Since then, PHAs have been detected in diverse bacteria (both autotrophs and heterotrophs), archaea (Poli et al., 2011), and some yeasts (Thu et al., 2023), where they typically accumulate under nutrient-limiting, carbon-rich conditions. PHB biosynthesis involves a core pathway encoded by *phaA* (β-ketothiolase), *phaB* (acetoacetyl-CoA reductase), and *phaC* (PHA synthase), which polymerizes 3-hydroxybutyryl-CoA. Additional regulators such as *phaR* and granule-associated *phaP* (phasin) influence polymer accumulation and granule morphology (Pötter et al., 2005). PHA synthases (PhaC enzymes) are grouped into four classes (I–IV) based on their substrate specificity and subunit composition, rather than their evolutionary lineage. Classes I, III, and IV primarily synthesize short-chain-length PHAs like PHB but differ in enzyme architecture: Class I enzymes are homodimers composed of two identical PhaC subunits; Class III enzymes function as heterodimers consisting of PhaC and a partner protein, PhaE; and Class IV enzymes also form heterodimers, pairing PhaC with PhaR. In contrast, Class II enzymes are specialized for producing medium-chain-length PHAs and typically include multiple PhaC isoforms (e.g., PhaC1 and PhaC2), without accessory subunits like PhaE or PhaR. A deeper understanding of how feedstock composition and PHA synthase structure influence polymer diversity and accumulation remains a key research goal.

*Cupriavidus necator* H16 is a model PHA producer and has been extensively engineered for high PHB yields (Zhang et al., 2022). Strategies include promoter and pathway optimization (Tang et al., 2020), substrate engineering (Mifune et al., 2010), and process development (Atlić et al., 2011). However, a key limitation is that PHB production in *C. necator* is not growth-associated, requiring nutrient shifts or batch processes, which hinders continuous cultivation (Henderson & Collins, 1997) and thus loss in industrial efficiency. Broader surveys have revealed PHA production in extremophiles, anaerobes, and oligotrophs (Wang et al., 2019a; Bennett et al., 2024), suggesting that novel chassis organisms with more favorable production dynamics remain to be explored.

Dissimilatory perchlorate-reducing bacteria (DPRB) represent a promising but understudied group in this context. Perchlorate (ClO_4_^−^) is a highly oxidized, water-soluble oxyanion with a biological redox potential comparable to oxygen (E°′ = +0.797 V) (Youngblut et al., 2016). In DPRB, perchlorate is reduced to chlorite by the perchlorate reductase complex (PcrAB), followed by dismutation into chloride and molecular oxygen via chlorite dismutase (Cld). The oxygen is subsequently respired by the microorganism (Youngblut et al., 2016). This pathway enables the endogenous generation of O_2_ under anoxic conditions, supporting oxygenase-dependent metabolism without external aeration (Carlström et al., 2015). The conserved, mobile perchlorate reduction island (PRI) allows this metabolism to be transferred across diverse taxa, supporting bioengineering efforts (Melnyk et al., 2015). *Azospira suillum* PS is a facultative anaerobe and genetically tractable DPRB capable of coupling perchlorate reduction to energy conservation and growth. Despite its unique metabolism and flexible redox physiology, its potential for bioproduct synthesis remains unexplored.

In this study, we investigate the genetic and physiological basis of PHB biosynthesis in *A. suillum* PS. Using comparative genomics, gene deletion, redox analysis, and polymer quantification, we characterize the pathways supporting PHB production during both aerobic and perchlorate-respiring growth. Our findings highlight *A. suillum* PS as a unique candidate for sustainable, growth-coupled bioplastic production in anaerobic biotechnological systems.

## Materials and Methods

### Bacterial Strains and Plasmids

*Azospira suillum* PS (ATCC BAA-33/DSMZ 13638) was revived from laboratory freezer stocks and used as the wild-type strain for all genetic manipulations. *Escherichia coli* XL1-Blue was used for plasmid propagation and cloning. A full list of primers (Supplementary Table 1), strains (Supplementary Table 2), and plasmids (Supplementary Table 3) is provided in the supplementary materials. Prior to all experiments, strains were streaked from freezer stocks to obtain single colonies.

### Mutant Construction

Gene deletion mutants in *A. suillum* PS were generated using a sacB-based counterselection approach (Gay et al., 1983). Electrocompetent cells were transformed with either integrative (∼500 ng) or replicative (∼50 ng) plasmids in 1 mm electroporation cuvettes using a Gene Pulser Xcell electroporator (Bio-Rad, USA) with the following settings: 1.8 kV, 25 µF, and 200 Ω. Cells were chilled on ice before electroporation; cuvettes were kept at room temperature. After pulsing, cells were recovered in pre-warmed ALP medium at 37°C for 6 h, then 100 µL of culture was plated directly on ALP agar containing 50 µg/mL kanamycin. Colonies appearing within 48 h were considered valid transformants and picked into 500 µL ALP broth with 50 µg/mL kanamycin. After 4 h of growth, cultures were plated on ALP agar with 6% sucrose for counterselection. Representative colonies were screened by colony PCR to confirm deletions. Glycerol stocks were prepared at 15% final concentration for storage at −80°C.

### Culture Conditions and Growth Media

*E. coli* strains were cultured in LB medium with 50 µg/mL kanamycin. Wild-type and mutant *A. suillum* PS strains were grown in acetate minimal medium (AMM). One liter of AMM contained: 0.49 g sodium phosphate monobasic dihydrate, 0.97 g sodium phosphate dibasic anhydrous, 0.1 g KCl, 0.25 g NH4Cl, and 10 mL each of vitamin and mineral mix (Bruce et al., 1999). The sodium acetate concentration in AMM was either 0.82g/L acetate for 10mM AMM or 2.05 g acetate for 25mM AMM. To prepare mixed carbon source media (acetate, lactate, pyruvate), AMM was supplemented with 2.0 g yeast extract, 7.6 mL of a 60% (w/w) sodium lactate solution, and 1.10 g sodium pyruvate. For solid media, 15 g/L agar was added. *A. suillum* PS was cultured in 250 mL baffled Erlenmeyer flasks containing 50 mL of medium at 37°C and 250 rpm. Optical density at 600 nm (OD_600_) was measured using a 1 mm pathlength cuvette on a Genesys 20 spectrophotometer (Thermo Scientific, USA).

### PHA Extraction and GC-MS Analysis

PHB was quantified from 40 mL of culture. Samples were harvested by centrifugation at 7,197 × g for 10 min at 4°C, and pellets were stored at −80°C. Acidic methanolysis was performed as described by Juengert et al. (2018). Extracts were analyzed by GC-MS using a 7890A GC system (Agilent Technologies, USA) equipped with a DB-WAX UI column and coupled to a 5975C XL EI/CI Mass Selective Detector. The temperature program was: 80°C for 2 min, ramp to 210°C at 10°C/min, then to 250°C at 50°C/min with a 1 min hold. Poly[(R)-3-hydroxybutyric acid] (Santa Cruz Biotechnology, USA) was used for the standard curve, and methyl benzoate (Sigma-Aldrich, USA) served as the internal standard. Compound identity was confirmed by total ion count (TIC) spectra matching the NIST EI database. The presence of polyhydroxyvalerate (PHV) and polyhydroxyhexanoate PHHX was assessed using a commercial PHA standard (Sigma-Aldrich, USA), and TIC peaks were compared with NIST spectra for 3-hydroxypentanoic acid methyl ester and 3-hydroxyhexanoic acid methyl ester.

### NAD(P)^+^/H Quantification

Redox metabolite analysis was performed as described by Kern et al. (2014). Samples were kept on ice and protected from light until incubated at 37°C. To prevent redox degradation, only three samples were processed at a time. Quantification was performed using a Varian Cary 50 MPR Microplate Reader (Agilent Technologies, USA).

### Fluorescence Microscopy

Microscopy was conducted as described by Juengert et al. (2018). Cells were harvested in exponential phase, washed once in PBS (8 g NaCl, 201 mg KCl, 1.42 g Na2HPO4, 245 mg KH2PO4 per liter, pH 7.4), and stained with 1 µg/mL Nile Red for 10 min in the dark. Samples were mounted on 1% agarose pads and visualized on a Zeiss LSM880 confocal microscope (Zeiss, Germany) using a 100× Plan-Apochromat NA 1.4 objective. Nile Red was excited at 561 nm, and emission was detected from 570–620 nm using a GaAsP photon-counting detector. Differential interference contrast (DIC) images were acquired simultaneously. Image acquisition settings were consistent across all samples and processed in ZEN software v2.3 SP1 (Zeiss).

### Phylogenetic Tree Construction

Each *phaC* gene from *A. suillum* PS was queried against the NCBI nr database using BLASTP (Altschul et al., 1990). The top 250 unique hits (excluding “multispecies” entries) were retained per gene. PhaC homologs from *Azospira* species were manually reviewed and validated using HMMER v3.4 with TIGRfam-specific models (Finn et al., 2011). Alignments were generated with MUSCLE v3.8.31 and trimmed using trimAl v1.4 (Edgar, 2004; Capella-Gutiérrez et al., 2009). Phylogenies were inferred using IQ-TREE v2.0.6 with 1,000 ultrafast bootstraps and the Q.pfam+I+R7 model. Trees were visualized using iTOL v6 (Minh et al., 2020; Letunic and Bork, 2020).

### In Silico Comparative Genomics

The genome of *A. suillum* PS was downloaded from NCBI (Sayers et al., 2024); *Dechloromonas agitata* CKB and additional *Azospira* genomes were retrieved from JGI-IMG (Markowitz et al., 2006). Genomes were searched using HMMER v3.4 with a custom set of TIGRfams relevant to PHA metabolism (Supplementary Table 4). Presence/absence patterns of PHA biosynthesis genes were visualized using R v4.4.2 and RStudio v2024.12.0+467.

## Results

### Genomic and microscopic evidence for PHB granule formation in *Azospira suillum* PS and other perchlorate-reducing bacteria

Previous work on the perchlorate-reducing bacterium *Dechloromonas agitata* CKB revealed electron-dense intracellular structures that were hypothesized to be PHA granules (Fig. 1a) (Bruce et al., 1999). *In silico* analysis (Fig. 1) confirmed that *D. agitata* CKB possesses all the core genes necessary for PHA biosynthesis, including *phaA, phaB, phaC, phaP*, and *phaR*, which together enable the conversion of acetyl-CoA to PHB and the regulation and granule-associated assembly of PHA. To further investigate this capacity in a more genetically tractable system, we turned to *Azospira suillum* PS, a model perchlorate-reducing bacterium with established genetic tools (Melnyk et al., 2014). Strain PS also contains the requisite genes for PHA biosynthesis (Fig. 1); however, unlike CKB, it additionally encodes the PHA depolymerase *phaZ*. Both organisms possess multiple copies of the *phaC* gene—three in CKB and four in PS (see Supplementary Table 5 for locus tags in PS). In both strains, one *phaC* copy is co-localized with *phaB* and *phaR* (Fig. 1), suggesting an operon structure, while the remaining *phaC* copies are dispersed throughout the genome. Of these, only one has a neighboring *phaP* gene, and no other adjacent genes associated with PHA biosynthesis were observed.

**Figure 1.**
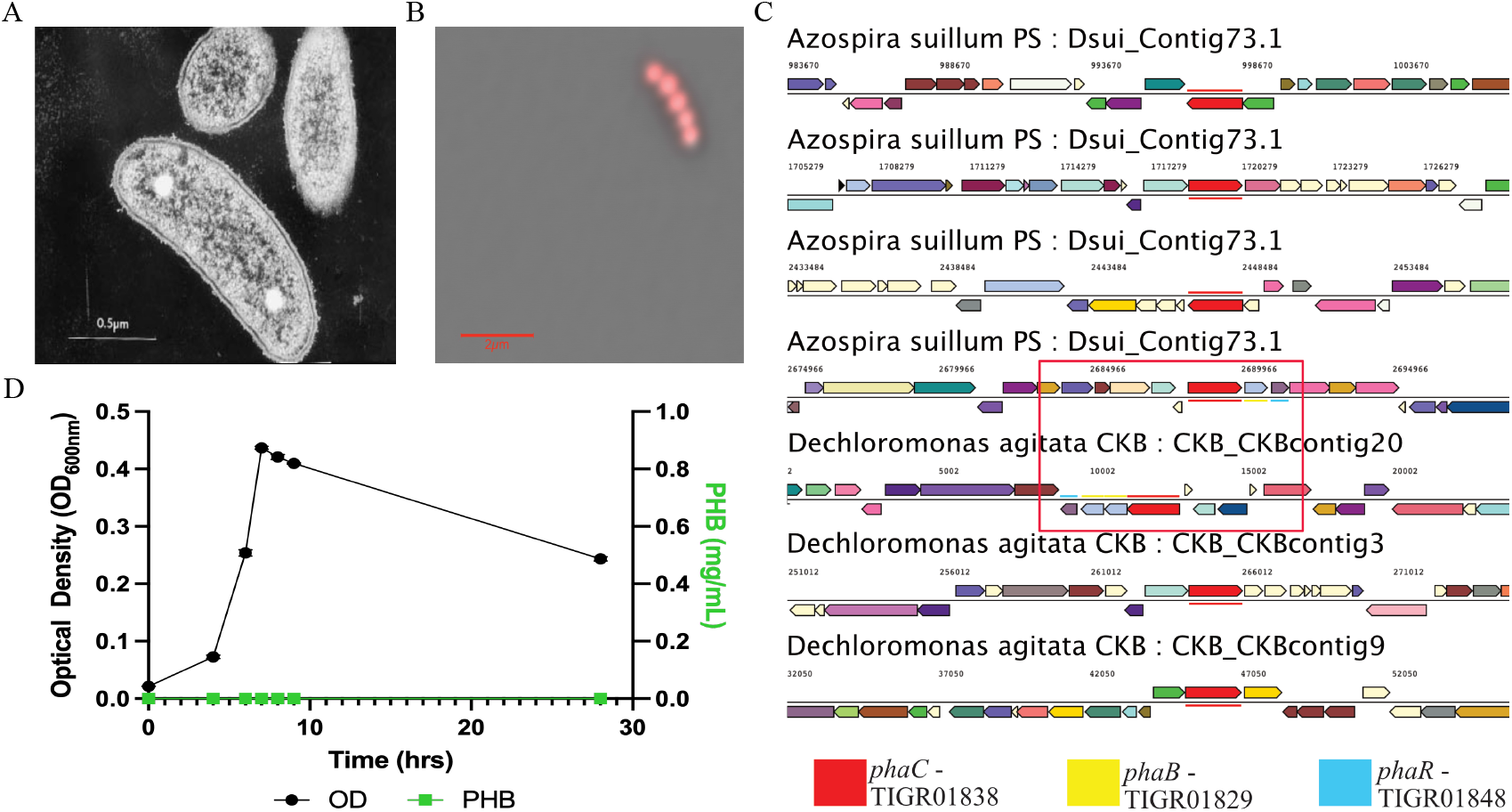
Microscopic and comparative genomic analysis of PHA production in *Dechloromonas agitata* CKB and *Azospira suillum* PS. (**A**) Transmission electron micrograph (TEM) of *D. agitata* CKB adapted from Bruce et al., 1995, showing electron-dense inclusions consistent with PHA granules. (**B**) Confocal fluorescence microscopy image of Nile Red-stained A. suillum PS overlaid with transmitted light, highlighting intracellular PHA granules. (**C**) Gene neighborhood analysis of phaC loci in CKB and PS, with neighboring PHA biosynthesis genes highlighted where present in the red box. (**D**) Growth (OD_600_) and absolute PHB production (mg/mL) by *A. suillum* PS during aerobic cultivation under nitrogen replete growth conditions.

To determine whether *A. suillum* PS produced PHB granules, we stained cultured cells with Nile Red, a lipophilic dye commonly used to visualize PHB granules and examined them using confocal microscopy (Fig. 1b). PS cells exhibited visible intracellular PHB granules with an approximate diameter of 0.32 µm. Given the ∼2 µm cell diameter of PS, the clustered granules span much of the intracellular space, corresponding to ∼64% of the estimated cell volume being occupied by PHB granules. Cells stained positive for PHB granules under both aerobic and perchlorate-respiring conditions, providing the first clear evidence of PHA production linked to perchlorate respiration. This observation confirmed the genetic potential for PHB accumulation; however, it remained unclear whether polymer synthesis occurred constitutively or only in response to specific nutrient conditions.

To establish whether PHB accumulation in *A. suillum* PS is constitutive or nutrient-dependent, we first evaluated growth under nitrogen-replete conditions. Cultures were grown in standard acetate minimal medium (AMM) containing 10 mM sodium acetate and 4.68 mM NH4Cl, corresponding to a replete C:N ratio of approximately 1:0.23 relative to the canonical Redfield ratio (C:N = 1:0.15) (Redfield, 1934). Under these conditions, no detectable PHB was observed by GC–MS analysis (Fig. 1d), confirming that PHB synthesis does not occur under nitrogen-replete growth. The culture exhibited a specific growth rate (µ) of 0.60 h−^1^ and a doubling time of 1.16 h, indicating robust biomass formation despite the absence of polymer accumulation.

Given prior genomic and microscopic evidence suggesting PHB-producing potential, we next examined whether altering the carbon-to-nitrogen ratio could induce polymer synthesis. When the acetate concentration was increased to 25 mM while maintaining 4.68 mM NH4Cl (C:N= 1:0.094), PHB production was observed under both aerobic and perchlorate-respiring conditions (Fig. 2a–d). Under these carbon-enriched conditions, cell experience ∼38% less nitrogen per unit carbon than under the canonical Redfield ratio, experiencing moderate nitrogen limitation. Under these conditions *A. suillum* PS exhibited a slightly reduced specific growth rate (µ = 0.50 h^−1^; doubling time = 1.39 h) but accumulated measurable intracellular PHB. These results demonstrate that PHB synthesis in *A. suillum* PS is triggered by a shift toward carbon surplus relative to nitrogen, rather than being a constitutive feature of its metabolism.

**Figure 2.**
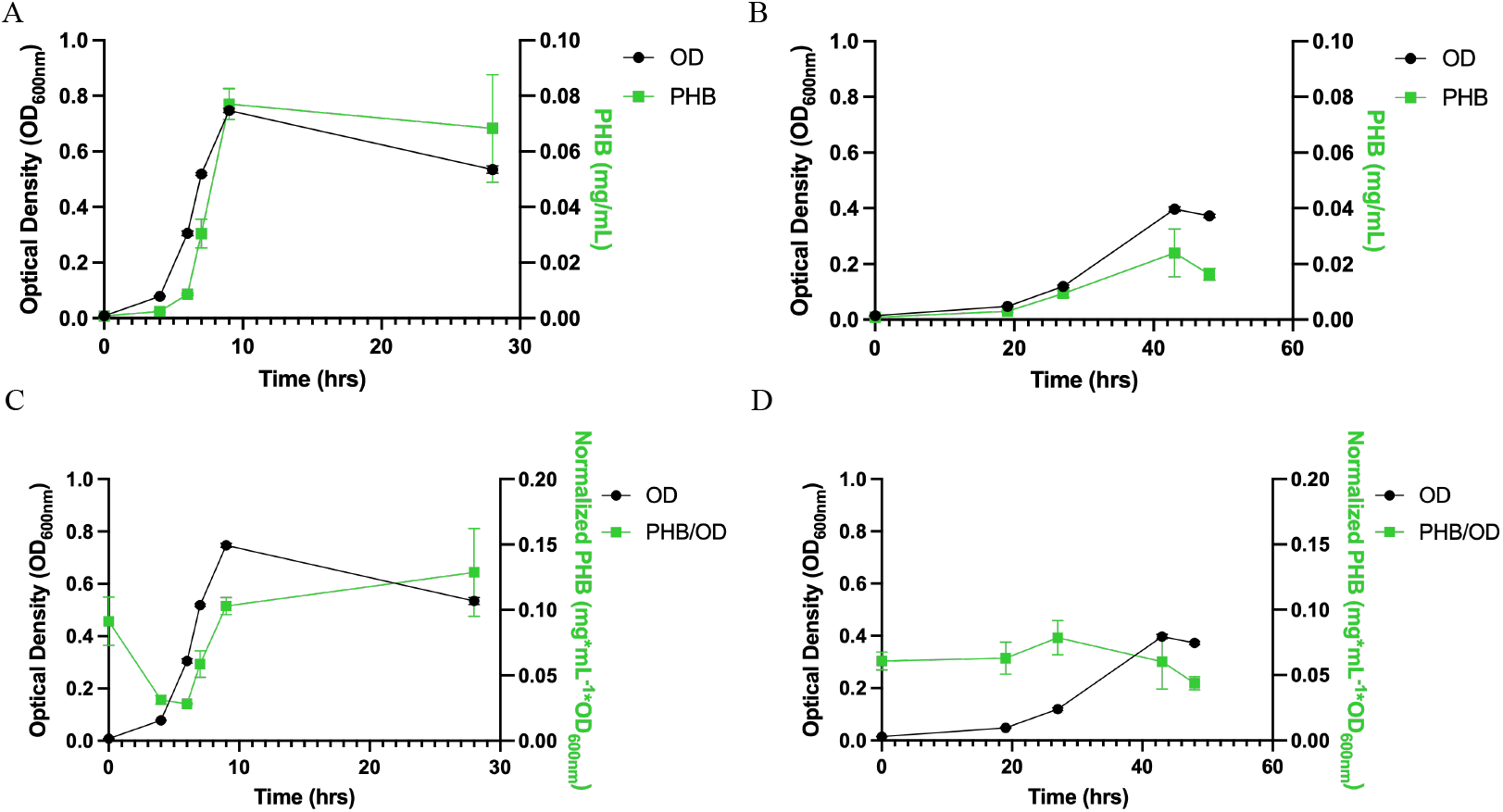
PHA production by *Azospira suillum* PS across electron acceptors with acetate as the electron donor under moderate nitrogen deplete conditions. **(A)** Growth (OD_600_) and absolute PHB production (mg/mL) during aerobic cultivation. **(B)** Growth (OD_600_) and absolute PHB production (mg/mL) during perchlorate-respiring growth. **(C)** Growth (OD_600_) and normalized PHB production (mg mL^−1^ OD_600_^−1^) under aerobic conditions. **(D)** Growth (OD_600_) and normalized PHB production (mg mL^−1^ OD_600_^−1^) during perchlorate respiration.

Following this finding, we conducted a detailed time-course analysis of PHB accumulation under the carbon-enriched condition to quantify the extent and dynamics of polymer synthesis during aerobic and perchlorate-respiring growth.

### Growth-associated PHB production in *Azospira suillum* PS under aerobic and perchlorate-respiring conditions

To quantify PHA production by *A. suillum* PS, gas chromatography–mass spectrometry (GC-MS) was performed on samples extracted via acidic methanolysis. In addition to quantifying total PHB produced by the culture, this method also enabled identification of the different monomers present. When PS was grown on acetate under either aerobic or perchlorate-respiring conditions 3-hydroxybutyric acid (3-HB) (the monomer of PHB) was detected. PHB was quantified across the growth curve and was found to be produced in a growth-associated manner (Fig 2a-d). While growth-associated PHB production has been previously reported (Scott et al., 2021; Amos & McInerney, 1989; Ackermann & Babel, 1997); PHB biosynthesis is more commonly observed during stationary phase, in response to nutrient limitation (Anderson & Dawes, 1990).

When grown aerobically, *A. suillum* PS reaches a maximum OD_600nm_ of 0.747 ± 0.008 during late exponential phase, at which point it had produced a maximum of 0.077 ± 0.006 mg/mL PHB providing a total yield of 0.10 g/L per OD_600nm_ (Fig 2a). Under perchlorate-respiring conditions with 10 mM perchlorate, PS reaches a lower maximum OD_600nm_ of 0.397 ± 0.009 and produces a maximum of 0.024 ± 0.008 mg/mL PHB or 0.06 g/L per OD_600nm_ (Fig 2b). When grown on perchlorate, PHB levels slightly decline as the culture enters stationary phase (Fig 2d). There is no significant reduction in PHB levels between peak and late-stage culture under aerobic conditions, despite a drop in OD_600nm_ of approximately 0.21. Based on the 1.6-fold higher PHB yield in aerobic versus perchlorate conditions, we further characterized PHB production by PS under aerobic growth.

This growth-associated pattern under carbon-enriched conditions contrasts with the nitrogen-replete culture, where no PHB was detected, reinforcing that polymer synthesis in *A. suillum* PS is activated only when the C:N ratio exceeds a physiological threshold. Under both mild and severe nitrogen limitation, PHB production remained unchanged, which is consistent with observations in other anaerobic organisms such as *Syntrophomonas wolfei*, which produced PHB independently of ammonium chloride concentration (Amos & McInerney, 1989).

When examining normalized PHB levels relative to optical density, both aerobic and perchlorate-grown cultures show evidence of carryover PHB from the inoculum (Fig 2c&d). In aerobic cultures, this residual PHB is rapidly degraded during lag phase prior to new PHB accumulation during log phase, suggesting that the cells may utilize the carryover PHB as an additional electron donor alongside the supplied acetate during the initial phase of growth (Fig 2c). In contrast, perchlorate-respiring cultures do not exhibit this initial degradation phase; instead, normalized PHB/OD_600_ remains stable during early growth and begins increasing only during exponential phase.

### Redox cofactor dynamics link NADPH availability to PHB biosynthesis in *Azospira suillum* PS

To assess the redox influence on PHB biosynthesis, we measured intracellular NADH/NAD^+^ and NADPH/NADP^+^ ratios over a 24-hour culture period and compared these values to PHB accumulation (Fig. 3a–d). When grown aerobically, *Azospira suillum* PS exhibited an average NADH/NAD^+^ ratio of 0.491—more than twice the reported aerobic values for *E. coli* (0.226) and *Klebsiella aerogenes* (0.195)—indicating a relatively reduced intracellular environment. Notably, this ratio also exceeded those observed under anaerobic conditions for both comparative organisms (0.396 and 0.219, respectively) (Wimpenny & Firth, 1972). Only *Pseudomonas aeruginosa*, a known respiratory generalist, exhibited a higher aerobic NADH/NAD^+^ ratio (0.571). While *E. coli* lacks the genetic capacity for native PHA production, some *Pseudomonas* spp. are known PHB producers, although not typically in a growth-associated fashion (Bertrand et al., 1990).

**Figure 3.**
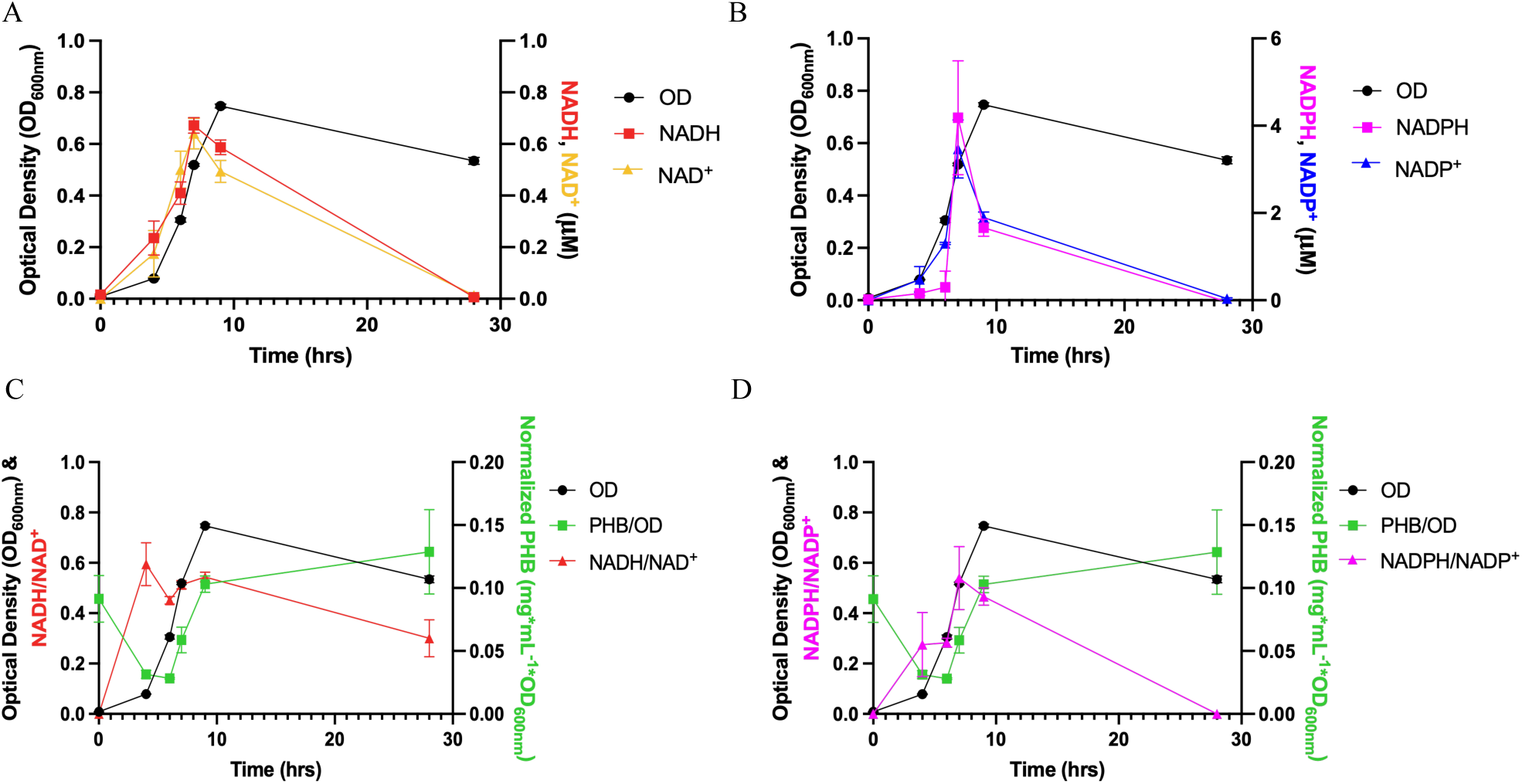
Normalized PHA production and intracellular redox state of *Azospira* suillum PS grown aerobically on acetate. **(A)** Growth (OD_600_) and absolute intracellular concentrations of NADH and NAD^+^ (µM). **(B)** Growth and absolute concentrations of NADPH and NADP^+^ (µM). **(C)** Growth, normalized PHB production (mg mL−^1^ OD_600_^−1^), and the NADH/NAD^+^ ratio. **(D)** Growth, normalized PHB production, and the NADPH/NADP^+^ ratio.

Spearman correlation analysis revealed a strong and statistically significant monotonic relationship between PHB levels and NADPH/NADP^+^ (ρ = 0.829, p = 0.042), while no significant correlation was observed with NADH/NAD^+^ (ρ = 0.600, p = 0.208). This is consistent with the known cofactor specificity of the PhaB acetoacetyl-CoA reductase enzyme. In *Cupriavidus necator* H16—a well-studied model organism for PHB production—this enzyme has been shown to preferentially use NADPH rather than NADH as the reducing equivalent source (Steinbüchel & Schlegel, 1991).

### Phylogenomic analysis reveals conserved, diversified *phaC* loci across the *Azospira* genus

A maximum-likelihood phylogenetic tree (Fig 4) constructed from the four unique *phaC* gene products identified in *Azospira suillum* PS, along with their 250 closest homologs, revealed that each PhaC protein is nested within a monophyletic clade composed of *phaC* homologs from other *Azospira* species. Despite this shared genus-level affiliation, the four *phaC* variants from strain PS occupy distinct and divergent branches within the tree, suggesting that each *phaC* gene may have evolved independently. Furthermore, this analysis revealed that multiple *phaC* gene copies are consistently present in *Azospira* genomes. Comparative genomic analysis across the genus showed that each *Azospira* species encodes at least three distinct copies of *phaC*, alongside a single copy of the transcriptional regulator *phaR*, suggesting that PHA biosynthesis is a conserved metabolic feature within the genus. To assess whether these genes may have been acquired via horizontal gene transfer, we used IslandViewer (Bertelli et al., 2017), which predicts genomic islands based on sequence composition and comparative genomics. This analysis showed no evidence of horizontal gene transfer near the *phaC* loci in strain PS (data not shown).

**Figure 4.**
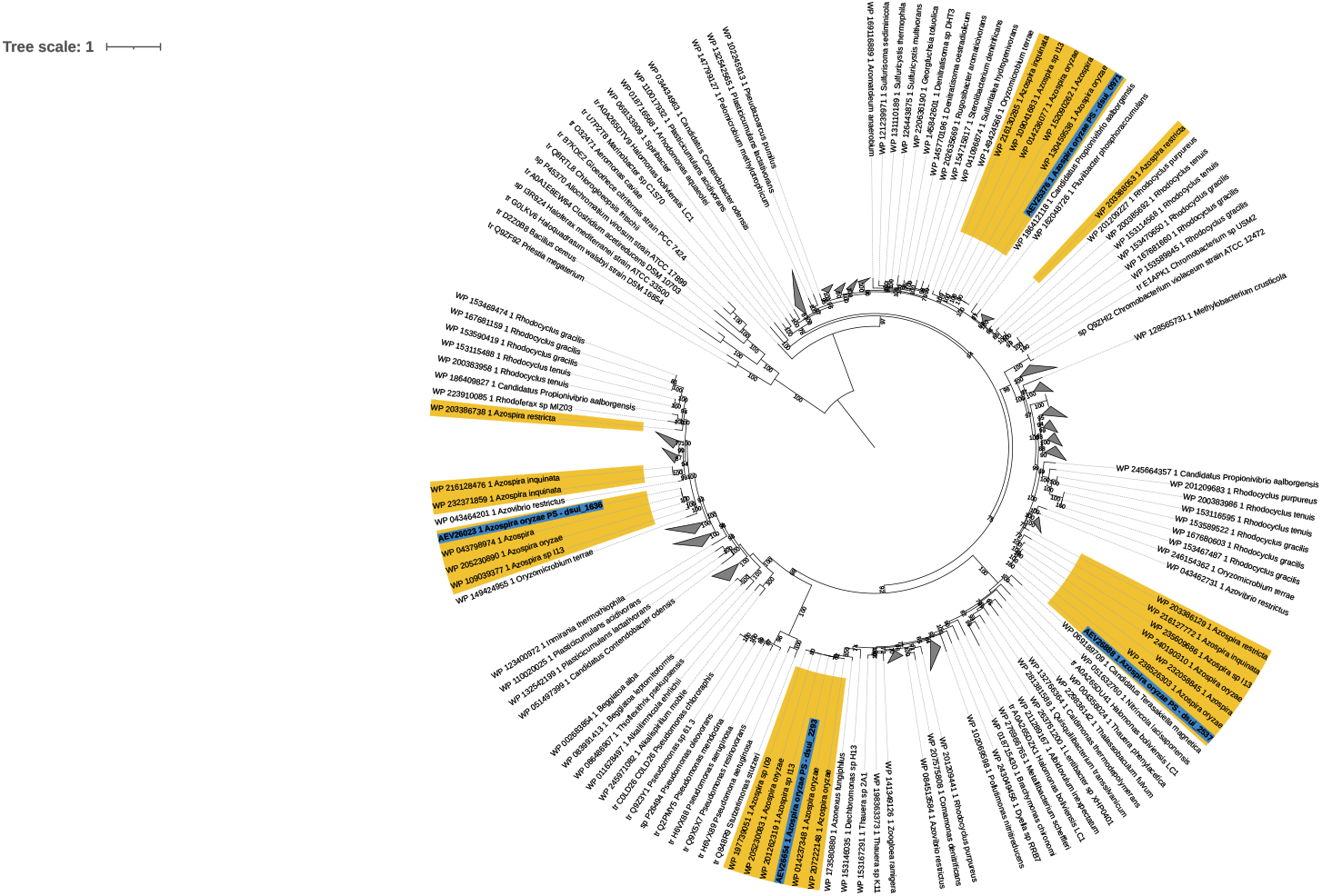
PhaC protein phylogeny within the genus *Azospira*. Maximum likelihood phylogenetic tree of PhaC protein sequences from members of the genus *Azospira*. Bootstrap support values (n = 1000 replicates) are shown at each node. Branches corresponding to *Azospira* species are highlighted in yellow, and those from A. suillum PS are further annotated in blue. Tip labels include the GenBank protein accession number followed by the corresponding NCBI organism name.

To assess conservation of gene neighborhoods, we used CAGE-CAT and Clinker to compare the genomic loci surrounding *phaC* across *Azospira* genomes (Gilchrist & Chooi, 2021). These tools align and visualize syntenic regions based on protein sequence identity and gene order, enabling rapid identification of conserved operons and structural rearrangements. This analysis (Fig 5) revealed that the local gene context of *phaC* is highly conserved, with pairwise amino acid identity of *phaC* homologs exceeding 70% across the genus, further supporting their functional importance in *Azospira*. However, while average nucleotide identity remains high across *Azospira* species, they do not share syntenic conservation of the broader PHA biosynthesis gene cluster.

**Figure 5.**
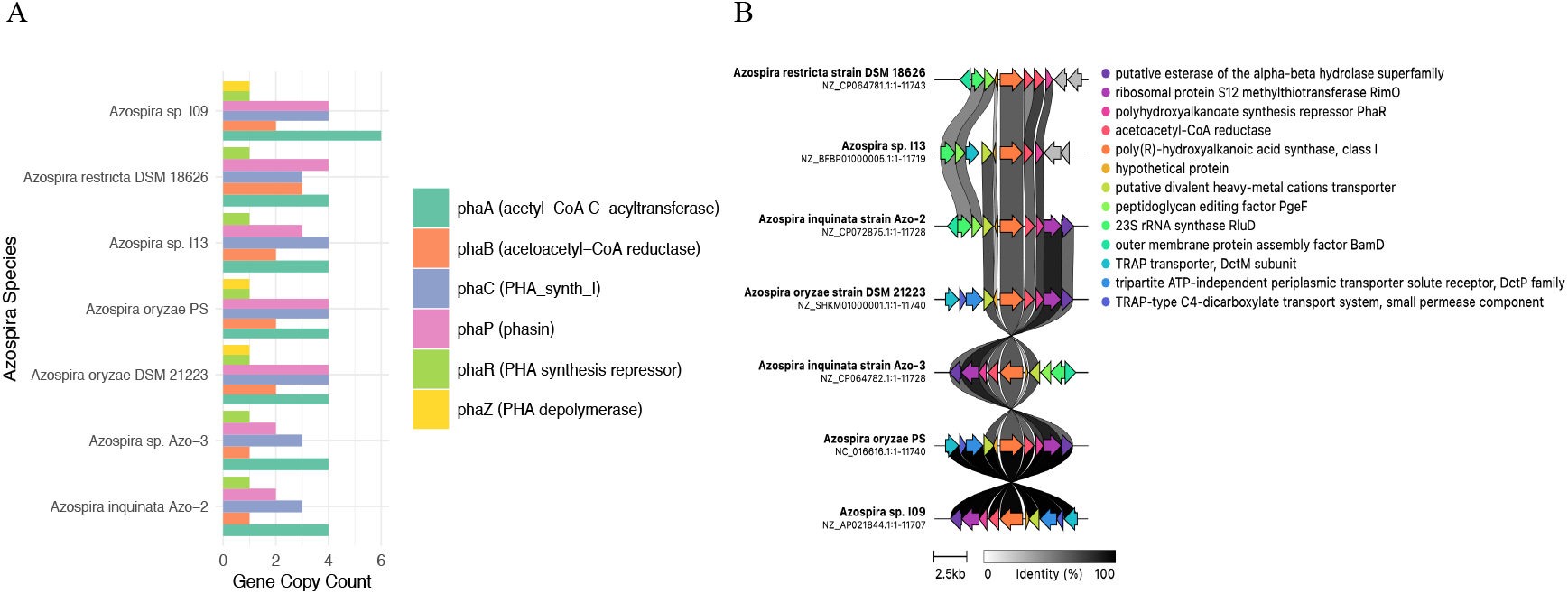
Distribution of PHA-associated genes across the genus *Azospira*. **(A)** Presence and distribution of all identified PHA-associated genes across sequenced genomes within the genus *Azospira*. **(B)** Nucleotide identity percentages for the phaC biosynthetic cluster co-localized with phaB and phaR across *Azospira* species, highlighting variation in conservation of this operon.

### Functional characterization of *phaC* genes reveals essential role of *phaC4* in PHB biosynthesis

Our analysis found that *Azospira suillum* PS encodes only class I *phaC* genes, which are typically associated with the production of short-chain-length (scl) PHAs such as PHB. To investigate the four *phaC* genes in *A. suillum* PS responsible for PHA production, we attempted to generate deletion mutants for each gene. Knockout strains were successfully constructed for three out of the four *phaC* genes. However, under all tested growth conditions, we were unable to generate a mutant lacking the *phaC* gene with the locus tag *dsui_2537* (*phaC*4). Notably, this gene is co-localized with *phaB* and *phaR*. Previous transposon mutagenesis studies in strain PS also identified *phaC*4 as essential under all conditions tested (Supplementary Figure 1).

To further assess the functional role of each *phaC*, we compared the PHA monomer profiles of all knockout strains under both defined and complex media conditions. As shown in Table 1, none of the knockout strains displayed a PHA profile distinct from wild type. All strains consistently produced PHB, while polyhydroxyvalerate (PHV) and polyhydroxyhexanoate (PHHX) were absent (data not shown). These results suggest that the essential *phaC*4 is playing an important physiological role during normal growth in nutrient replete media.

**Table 1.**
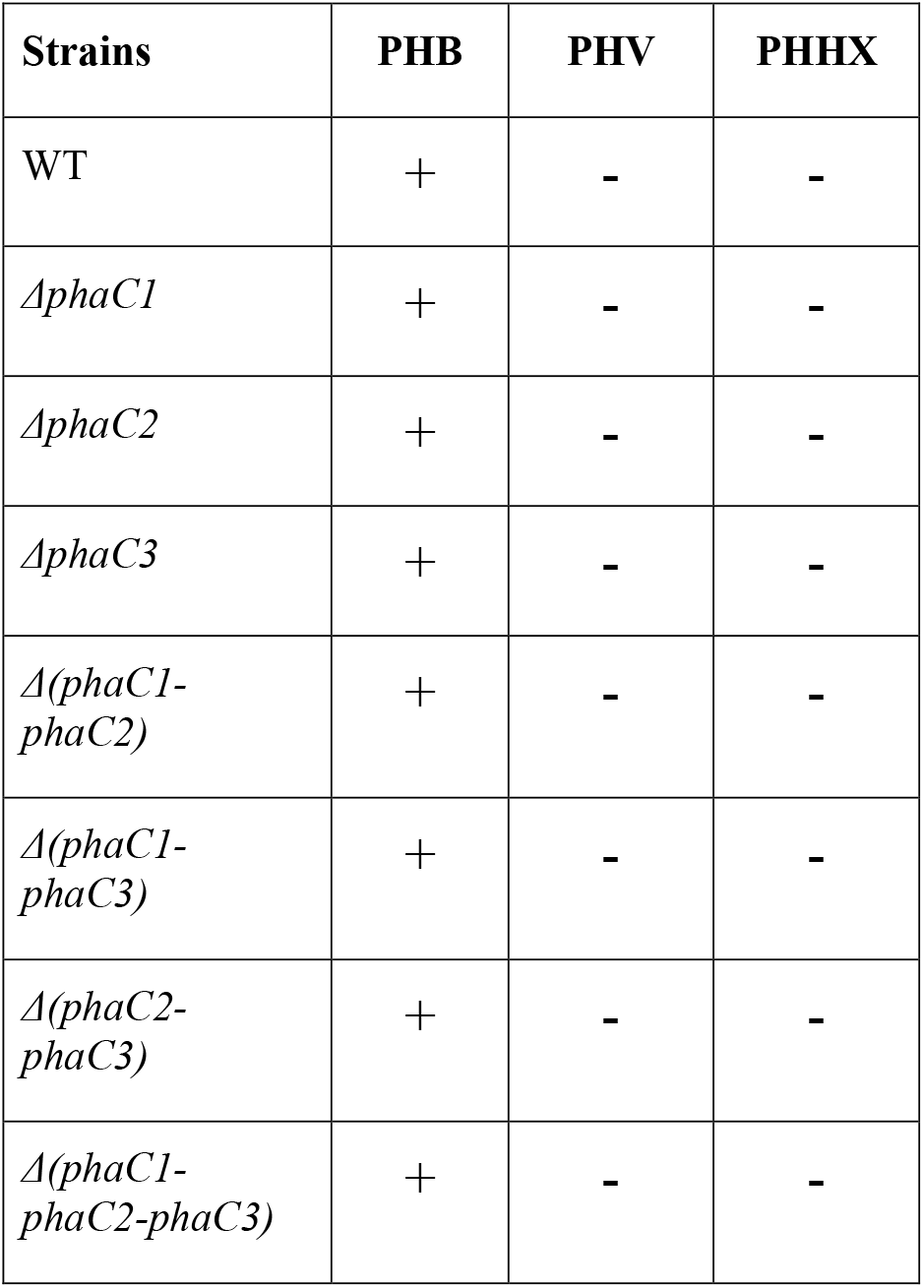
Polyhydroxyalkanoate methyl esters produced by ***Azospira suillum*** PS when grown on complex or defined media. Summary of PHA monomers identified in *A. suillum* PS when cultivated on complex media (acetate, lactate, pyruvate with yeast extract) or defined acetate minimal medium. PS locus tags corresponding to each *phaC* gene are listed in Supplementary Table 5.

## Discussion

This study provides new insight into the biosynthesis and regulation of polyhydroxybutyrate (PHB) in *Azospira suillum* PS, a model perchlorate-reducing bacterium. We demonstrate for the first time that PHB biosynthesis occurs under perchlorate-respiring conditions, expanding the known metabolic range for polyhydroxyalkanoate (PHA) production. Notably, *A. suillum* PS accumulates PHB in a growth-associated manner under both aerobic and anaerobic conditions, with aerobic cultures producing 1.6-fold more PHB. PHB synthesis was absent under nitrogen-replete conditions and was only observed when the carbon-to-nitrogen ratio was altered to a ∼40 % reduction in nitrogen availability relative to the canonical Redfield ratio. This establishes that PHB production in *A. suillum* PS is triggered by moderate nutrient imbalance. This pattern contrasts with the stress-induced, stationary-phase accumulation commonly observed in other bacteria, such as *C. necator*, suggesting that PHB plays a role in the central metabolism of *A. suillum* PS under nitrogen limitation where it is coupled to growth under both aerobic and perchlorate reducing conditions.

Temporal analysis of PHB accumulation revealed distinct regulatory patterns between respiratory modes. Under aerobic growth, normalized PHB/OD_600_ initially declines during lag phase, likely reflecting mobilization of stored carbon and reducing equivalents. In contrast, perchlorate-respiring cultures do not exhibit this early decrease; instead, PHB levels remain stable in early growth and begin to rise during exponential phase.

Our redox profiling data support this hypothesis. A strong and statistically significant correlation was observed between intracellular NADPH/NADP^+^ ratios and PHB content (Spearman ρ = 0.829, *p* = 0.042), implicating NADPH availability as a potential driver of PHB accumulation. Although this observation suggests a role for NADPH in fueling PHB biosynthesis, this correlation does not confirm cofactor specificity of the biosynthetic enzymes. NADH and NADPH levels are often tightly coupled and metabolically interconvertible, and the observed association may reflect a broader redox state rather than direct NADPH usage. Further biochemical characterization will be required to determine whether PhaB or related enzymes in *A. suillum* PS preferentially utilize NADPH over NADH.

Genomic analysis revealed four *phaC* genes encoding class I PHA synthases in *A. suillum* PS. Of these, *phaC4* is particularly notable for its synteny with *phaB* and *phaR*, suggesting a conserved operon structure. No evidence of horizontal gene transfer was detected near any *phaC* loci, indicating that these genes are native and vertically inherited. Functional analysis supports the essentiality of *phaC4*, as multiple attempts to generate a knockout mutant were unsuccessful under all tested conditions. These findings further reinforce the hypothesis that PHB biosynthesis plays a central physiological role in this organism, potentially as a buffer for intracellular reducing equivalents.

From an applied perspective, *A. suillum* PS demonstrates rapid and efficient PHB accumulation on simple substrates. Under aerobic conditions, wild-type PS produces 8.5 mg/L/h PHB, reaching 77 mg/L in just 9 hours using a single 25 mM acetate dose (600 mg/L carbon). While this yield is lower than that of *Cupriavidus necator*—which reaches up to 79.2 mg/L/h on soybean oil after 72 hours—*C. necator* requires multiple feedings totaling ∼7,000 mg/L carbon (Mifune et al., 2010), greater than tenfold more than the PS culture. In contrast, PS grows in minimal medium without supplementation, avoiding the cost and handling challenges of plant oils. These characteristics position *A. suillum* PS as a promising chassis for continuous, low-input bioplastic production.

With targeted metabolic engineering—such as overexpression of *phaCBR* and optimization of carbon feed strategies—*A. suillum* PS may achieve higher titers while retaining its growth-associated PHB phenotype. This production strategy is particularly well-suited for bioprocesses favoring continuous synthesis and redox-balanced metabolism. Overall, our findings position *A. suillum* PS as a novel and sustainable chassis for industrial PHA production, especially in systems where simplicity, growth-coupling, and cost-effective carbon utilization are critical design goals.

### Industrial relevance

This study identifies *Azospira suillum* PS as a promising new chassis for sustainable bioplastic production. Unlike most known PHA producers, *A. suillum* PS accumulates PHB in a growth-associated manner under both aerobic and anaerobic (perchlorate-respiring) conditions, enabling simpler, continuous fermentation processes. Its ability to produce PHB rapidly on low-cost substrates in minimal media, combined with native genetic tractability and redox-linked regulation, makes it an attractive platform for scalable, low-input, and flexible biomanufacturing—particularly in resource-limited or oxygen-poor environments.

## Author Contributions

DAOM, HKC and JDC designed research. DAOM and BG optimized the genetic protocol and BG constructed the mutants. DAOM and VCL generated the phylogenetic analysis. DAOM performed all physiology experiments and measurements and comparative genomic analysis. DAOM wrote the draft manuscript and created the figures with guidance from JDC. All authors contributed to data analysis, reviewed the manuscript and approved of its publication.

## Acknowledgments

The authors acknowledge Yi Liu for guidance on instrumentation and analyses, Dr. Denise Schichnes at the UC Berkeley Biological Imaging Facility for training and access to microscopy, and Dr. Ruiwen Hu for his advice on computational analyses.

