## Supplementary Figures 1-3 for "Bioplastic Production Potential of *Azospira suillum* PS: Growth-Associated PHB Production Under Aerobic and Anaerobic Conditions": meier_et_al_supplementary_figures.pdf

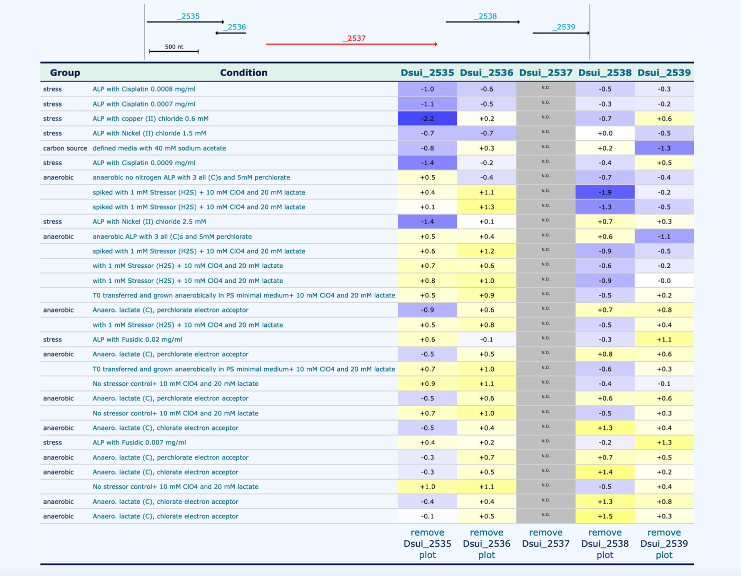


| Group | Condition | Dsui_2535 | Dsui_2536 | Dsui_2537 | Dsui_2538 | Dsui_2539 |
|---|---|---|---|---|---|---|
| stress | ALP with Cisplatin 0.0008 mg/ml | -1.0 | -0.6 | N.D. | -0.5 | -0.3 |
| stress | ALP with Cisplatin 0.0007 mg/ml | -1.1 | -0.5 | N.D. | -0.3 | -0.2 |
| stress | ALP with copper (II) chloride 0.6 mM | -2.2 | +0.2 | N.D. | -0.7 | +0.6 |
| stress | ALP with Nickel (II) chloride 1.5 mM | -0.7 | -0.7 | N.D. | +0.0 | -0.5 |
| carbon source | defined media with 40 mM sodium acetate | -0.8 | +0.3 | N.D. | +0.2 | -1.3 |
| stress | ALP with Cisplatin 0.0009 mg/ml | -1.4 | -0.2 | N.D. | -0.4 | +0.5 |
| anaerobic | anaerobic no nitrogen ALP with 3 all (C)s and 5mM perchlorate | +0.5 | -0.4 | N.D. | -0.7 | -0.4 |
| | spiked with 1 mM Stressor (H2S) + 10 mM ClO4 and 20 mM lactate | +0.4 | +1.1 | N.D. | -1.9 | -0.2 |
| | spiked with 1 mM Stressor (H2S) + 10 mM ClO4 and 20 mM lactate | +0.1 | +1.3 | N.D. | -1.3 | -0.5 |
| stress | ALP with Nickel (II) chloride 2.5 mM | -1.4 | +0.1 | N.D. | +0.7 | +0.3 |
| anaerobic | anaerobic ALP with 3 all (C)s and 5mM perchlorate | +0.5 | +0.4 | N.D. | +0.6 | -1.1 |
| | spiked with 1 mM Stressor (H2S) + 10 mM ClO4 and 20 mM lactate | +0.6 | +1.2 | N.D. | -0.9 | -0.5 |
| | with 1 mM Stressor (H2S) + 10 mM ClO4 and 20 mM lactate | +0.7 | +0.6 | N.D. | -0.6 | -0.2 |
| | with 1 mM Stressor (H2S) + 10 mM ClO4 and 20 mM lactate | +0.8 | +1.0 | N.D. | -0.9 | -0.0 |
| | T0 transferred and grown anaerobically in PS minimal medium+ 10 mM ClO4 and 20 mM lactate | +0.5 | +0.9 | N.D. | -0.5 | +0.2 |
| anaerobic | Anaero. lactate (C), perchlorate electron acceptor | -0.9 | +0.6 | N.D. | +0.7 | +0.8 |
| | with 1 mM Stressor (H2S) + 10 mM ClO4 and 20 mM lactate | +0.5 | +0.8 | N.D. | -0.5 | +0.4 |
| stress | ALP with Fusidic 0.02 mg/ml | +0.6 | -0.1 | N.D. | -0.3 | +1.1 |
| anaerobic | Anaero. lactate (C), perchlorate electron acceptor | -0.5 | +0.5 | N.D. | +0.8 | +0.6 |
| | T0 transferred and grown anaerobically in PS minimal medium+ 10 mM ClO4 and 20 mM lactate | +0.7 | +1.0 | N.D. | -0.6 | +0.3 |
| | No stressor control+ 10 mM ClO4 and 20 mM lactate | +0.9 | +1.1 | N.D. | -0.4 | -0.1 |
| anaerobic | Anaero. lactate (C), perchlorate electron acceptor | -0.5 | +0.6 | N.D. | +0.6 | +0.6 |
| | No stressor control+ 10 mM ClO4 and 20 mM lactate | +0.7 | +1.0 | N.D. | -0.5 | +0.3 |
| anaerobic | Anaero. lactate (C), chlorate electron acceptor | -0.5 | +0.4 | N.D. | +1.3 | +0.4 |
| stress | ALP with Fusidic 0.007 mg/ml | +0.4 | +0.2 | N.D. | -0.2 | +1.3 |
| anaerobic | Anaero. lactate (C), perchlorate electron acceptor | -0.3 | +0.7 | N.D. | +0.7 | +0.5 |
| anaerobic | Anaero. lactate (C), chlorate electron acceptor | -0.3 | +0.5 | N.D. | +1.4 | +0.2 |
| | No stressor control+ 10 mM ClO4 and 20 mM lactate | +1.0 | +1.1 | N.D. | -0.5 | +0.4 |
| anaerobic | Anaero. lactate (C), chlorate electron acceptor | -0.4 | +0.4 | N.D. | +1.3 | +0.8 |
| anaerobic | Anaero. lactate (C), chlorate electron acceptor | -0.1 | +0.5 | N.D. | +1.5 | +0.3 |
| | | remove Dsui_2535 plot | remove Dsui_2536 plot | remove Dsui_2537 | remove Dsui_2538 plot | remove Dsui_2539 plot |

**Supplementary Figure 1 |** Transposon sequencing barcode data showing fitness of insertion mutants in the *phaCBR* gene cluster of *Azospira suillum* PS. The essentiality of *phaC* (dsui_2537) is supported by the absence of viable transposon insertions across growth conditions.

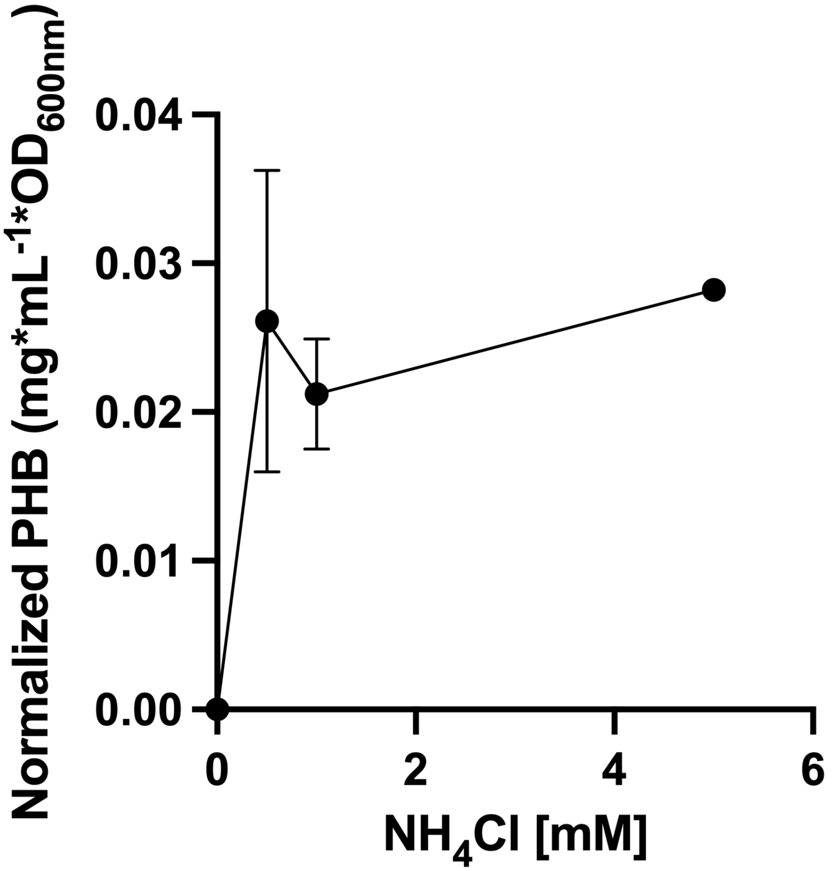


**Supplementary Figure 2 |** Effect of ammonium chloride concentration on normalized PHB production by *Azospira suillum* PS grown aerobically on 25 mM acetate. Samples were collected during early exponential phase.

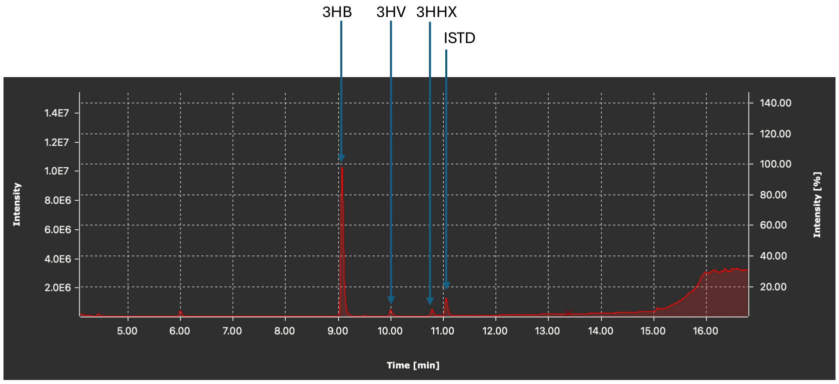


**Supplementary Figure 3 |** GC-MS spectra of a commercial mixed PHA standard, confirming the presence of all tested monomers from a validated source.

**Supplementary Table**

**Table S1 |** Primer List

**Table S2 |** Strain List

**Table S3 |** Plasmid List

**Table S4 |** PHA TIGRfams

**Table S5 |** PS PHA Locus Tags
